# High-resolution analysis of spatiotemporal virulence gene regulation during food-borne infection with *Escherichia coli* O157:H7 within a live host

**DOI:** 10.1101/128934

**Authors:** Daniel H Stones, Alexander GJ Fehr, Thekke P Madhavan, Kerstin Voelz, Anne Marie Krachler

## Abstract

Food-borne infection with enterohemorrhagic *Escherichia coli* (EHEC) is a major cause of diarrheal illness in humans, and can lead to severe complications such as hemolytic uremic syndrome. Cattle and other ruminants are the main reservoir of EHEC, which enters the food-chain through contaminated meat, dairy, or vegetables. However, how EHEC transitions from the transmission vector to colonizing the intestinal tract, and how virulence-specific genes are regulated during this transition, is not well understood. Here, we describe the establishment of a vertebrate model for food-borne EHEC infection, using the protozoan *Paramecium caudatum* as a vector and the zebrafish (*Danio rerio*) as a host. At 4 days post fertilization, zebrafish have a fully developed intestinal tract, yet are fully transparent. This allows us to follow intestinal colonization, microbe-host cell interactions, and microbial gene induction within the live host and in real time throughout the infection. Additionally, this model can be adapted to compare food- and water-borne infections, under gnotobiotic conditions or against the backdrop of an endogenous (and variable) host microbiota. Finally, the zebrafish allows for investigation of factors affecting shedding and transmission of bacteria to naïve hosts. High-resolution analysis of EHEC gene expression within the zebrafish host emphasizes the need for tight transcriptional regulation of virulence factors for within-host fitness.

**IMPORTANCE:** Enterohemorrhagic *Escherichia coli* (EHEC) is a food-borne pathogen which can cause diarrhea, vomiting and in some cases, severe complications such as kidney problems in humans. Up to 30% of cattle are colonized with EHEC, which can enter the food-chain through contaminated meat, dairy and vegetables. In order to control infections and stop transmission, it is important to understand what factors allow EHEC to colonize its hosts, cause virulence and aid transmission. Since this cannot be systematically studied in humans, it is important to develop animal models of infection and transmission. We developed a model which allows us to study food-borne infection in zebrafish, a vertebrate host that is transparent and genetically tractable. Using the zebrafish host, we can follow the bacterial infection cycle in real time, and gain important information regarding bacterial physiology and microbe-host interactions. This will allow us to identify potential new targets for infection control and prevention.

## INTRODUCTION

Enterohemorrhagic *Escherichia coli* (EHEC) are a major cause of food-borne infections worldwide. EHEC are transmitted through consumption of water, meat, dairy or vegetables contaminated with fecal matter, or hand-to-mouth, which is common in school and nursery settings. EHEC infection usually presents with bloody diarrhea, vomiting and stomach cramps, but in rare cases can lead to hemolytic uremic syndrome (HUS), a severe clinical complication resulting in kidney damage and often life-long morbidity, or mortality. Antibiotics are contraindicated, since antibiotic treatment can increase toxin production and exacerbate toxin-mediated disease pathology. Depending on geographical location, up to 30 % of cattle are colonized by EHEC, which presents a considerable environmental reservoir. One of the main virulence factors associated with colonization of ruminants as well as human hosts is the locus of enterocyte effacement (LEE), a horizontally acquired pathogenicity island encoding for a type 3 secretion system (T3SS). The LEE also encodes for the adhesion intimin and its host translocated receptor Tir (translocated intimin receptor), which in volunteer studies with the closely related enteropathogenic *E. coli* (EPEC) have shown to be a key factor for the development of diarrheal symptoms (1).

Ongoing studies of EHEC focus on understanding how the LEE is regulated during the EHEC life cycle, and how the LEE encoded genes and other virulence factors, such as Shiga toxin (STx) contribute to colonization, disease pathogenesis and transmission. Another area of interest is how a host’s endogenous microbiota interacts with EHEC, and how this affects host fate following EHEC ingestion. Several infection models exist to study EHEC virulence factors, most notably pigs, rabbits and mice. And although no single model host is capable of reproducing the full clinical presentation of human EHEC infection, each has its own distinct advantages and disadvantages (2). Gnotobiotic piglets inoculated with EHEC present a model for GI tract pathology, but fail to develop systemic symptoms. Additionally, this model is expensive, requires dedicated and highly specialized facilities, and is not particularly genetically tractable. Inoculation of neonate or infant rabbits through gastric catheters leads to colonization, diarrhea and GI tract histopathology that resembles that of human infection, but no mortality (3-5). Mice present the least expensive and most widely available vertebrate infection model to date, but their endogenous microbiota prevents EHEC colonization and has to be removed by streptomycin treatment to allow infection. Streptomycin treated mice can be used to study colonization, but fail to develop diarrhea, colitis, or A/E lesions (6). All three models present limited opportunities to study phenotypical changes in microbe and host simultaneously, in real time, and over a prolonged period. Intravital imaging of intestinal infection in these vertebrate models is possible but challenging, since part of the intestine has to be surgically exposed, which constitutes a severe procedure and can only be done for a limited duration. Additionally, it is technically challenging to study host-microbe interactions at a single cell level in this context.

Here, we set out to study EHEC colonization, host-microbe interactions and transmission in zebrafish (*Danio rerio*), a vertebrate host that is inexpensive, gives rise to a large offspring with short generation times, is genetically tractable and transparent during infancy. Due to these features, zebrafish larvae have become a well-established model for many bacterial (7, 8), viral (9) and fungal (10) infections, and for pathogen-microbiota interactions (11, 12). Genetic tractability paired with its transparency mean that phenotypical features of both microbe and host, and their change in response to interaction, can be studied during infection within a live host, over extended periods, and in a high-throughput format (13, 14). Changes in microbial and host gene expression, immune cell recruitment, and changes in host tissue morphology and intracellular signaling thus all become directly observable (15).

We capitalized on these features to study the dynamics of colonization, pathogen-microbiota interaction, and the effect of previously described virulence factors on microbe-host interaction following food-borne infection of infant zebrafish with EHEC.

## RESULTS

### *Paramecium caudatum* acts as a vector for *Escherichia coli*

Existing animal models predominantly administer *E. coli* as liquid suspension, through oro-gavage (2). In contrast, infections in humans usually result from ingestion of contaminated food. The unicellular ciliated protozoan, *Paramecium caudatum*, is used as a natural food-source for zebrafish larvae. It is abundant in freshwater, brackish and marine water, and has been described as a reservoir for environmental bacteria (16) as well as a vector for fish diseases (17). Thus, we tested whether it could be established as a vector for food-borne infection of zebrafish larvae with EHEC.

*P. caudatum* preys on bacteria, algae or yeasts, which it takes up via its ciliated oral groove, delivers into its mouth opening, from where it reaches the gullet. Eventually, ingested material blebbs and is internalized into a vacuole, which gradually acidifies and renders contents subject to degradation. We studied the interactions between *P. caudatum* and the enterohemorrhagic *E. coli* O157:H7 Sakai strain, to establish if and for how long bacteria would persist within *Paramecium. P. caudatum* was able to utilize *E. coli* as food source, as paramecia proliferated when kept in co-culture, in a dose-dependent manner (Fig. 1A). Fluorescence imaging following co-culture with EHEC*:mCherry* revealed that paramecia contained *E. coli*, which were localized within food vacuoles (Fig. 1B-D). Ingestion of EHEC did not seem to have any adverse effects on *P. caudatum* (Fig. 1B) and indeed, paramecia were observed to proliferate on a diet of EHEC (Fig. 1C). Further investigation using co-culture of *P. caudatum* and EHEC:*gfp* revealed that *E. coli* containing food-vacuoles remain transiently stable, but eventually acidify (Fig. 1D), leading to degradation of *E. coli*. The time course of degradation was determined by plating of viable *E. coli* recovered from paramecium (Fig. 1E). Ingested *E. coli* were degraded with a half-life of approximately 2.5 hours (Fig. 1E). This aligns with our visual confirmation of EHEC degradation within paramecia (Fig. S1). Dilution plating of EHEC-containing paramecia grown on EHEC for two hours showed that each *P. caudatum* contained an average of 200 viable *E. coli*, and that the number of intracellular EHEC was independent of bacterial density over a concentration range of 2.5.10^7^ to 10^8^ CFU/mL. This allowed us to establish *P. caudatum* as a vector to administer a defined dose of food-borne EHEC to zebrafish larvae.

**Fig. 1.**
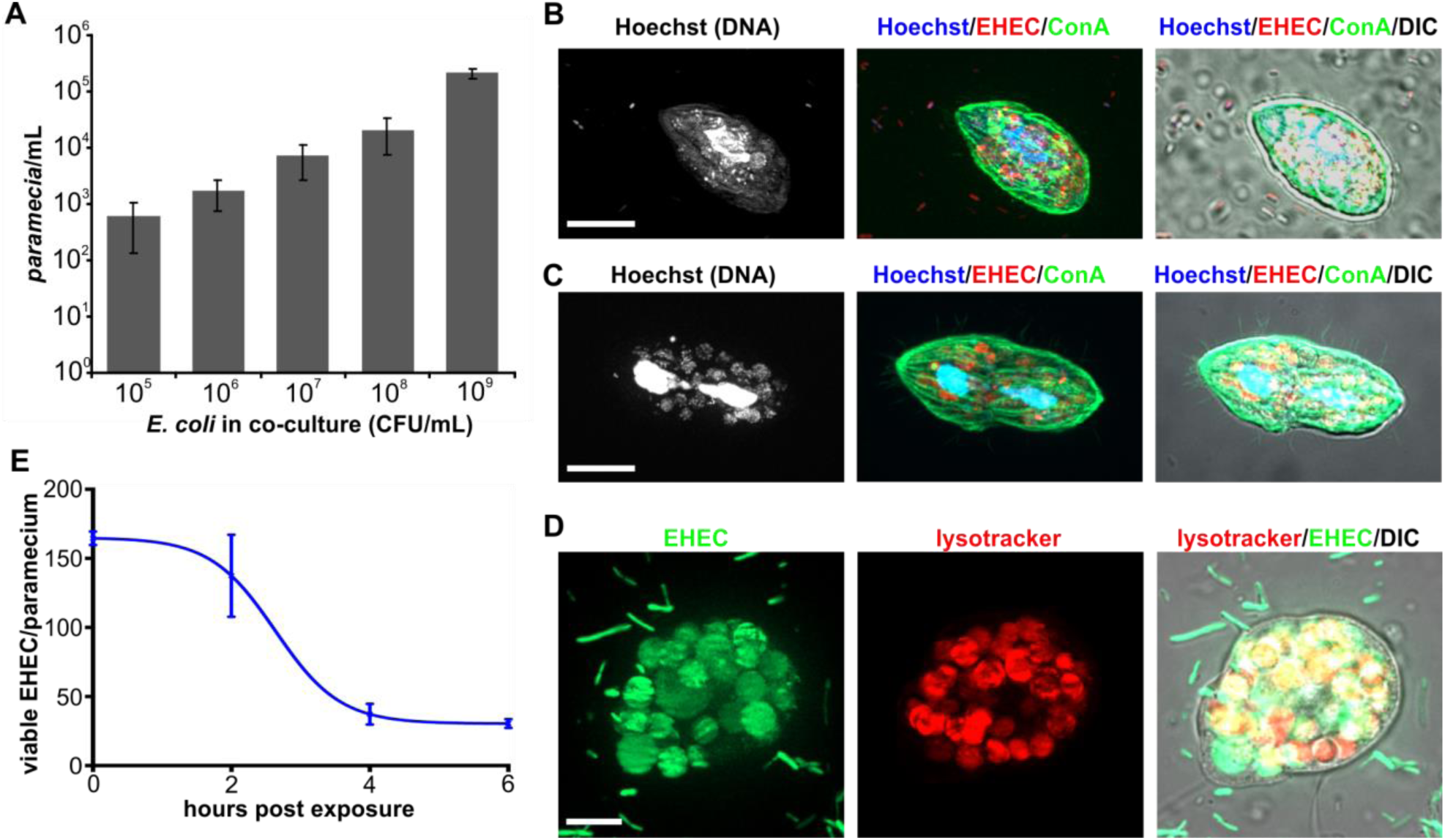
*Paramecium caudatum* acts as a vector for *Escherichia coli*. (A) EHEC and *P. caudatum* were grown in co-culture for 16 hours, and *P.caudatum* numbers measured using a hemocytometer. Numbers of paramecia at the end of the experiment were graphed against initial bacterial concentrations (CFU/ml). Values shown are means ± stdev. (n=3). (B) Following 2 hours of co-incubation with EHEC:mCherry (red), *P. caudatum* samples were fixed and stained with Hoechst (DNA, blue) and concanavalin A (green). Some paramecia were proliferating on a diet of EHEC, and undergoing cell division (C) Scale bars, 10 μm. (D) Following 2 hours of co-culture with EHEC:gfp (green), *P. caudatum* were incubated with lysotracker (red) to visualize vacuole acidification. Scale bar, 5 μm (E) Following two hours of co-incubation with EHEC, *P. caudatum* were transferred to medium without bacteria. Samples were removed at indicated timepoints and numbers of viable *E. coli* within paramecia were determined by dilution plating on selective agar (n=3).

### Food-borne *E. coli* colonize larval zebrafish more efficiently than water-borne EHEC

Initially, colonization by water-borne EHEC was characterized in gnotobiotic zebrafish larvae, which were acquired from bleached eggs and reared under sterile conditions as previouly described (18). EHEC were administered to larval zebrafish at 4 dpf, at which point larvae had a fully developed intestinal tract with a functional anal opening. *E. coli* constitutively expressing mCherry were used to visualize colonization *in vivo*. Bacterial burden was initially assessed following 2 hours of exposure, and increased dependent on the amount of ingested EHEC (Fig. 2A). Colonization was mostly localized to the mid-intestine (Fig. 2B), and accumulation and secretion of EHEC-containg fecal matter was observed soon after ingestion (Fig. 3).

Next, paramecia loaded with EHEC Sakai:mCherry were administered to larval zebrafish at 4 dpf, at which point larvae were able to swim freely, and prey on paramecium (Figure S2 and Movie S1). To administer a consistent number of *E. coli* to zebrafish, paramecia were kept in co-culture with *E. coli* at a ratio of 1:500, gently washed to remove extracellular *E. coli* and added to fish medium to result in a defined initial concentration as described above, and offered to zebrafish as a prey.

Bacterial burden was determined in fish exposed to matched doses of food-borne or water-borne EHEC, following two hours of exposure. At all three doses tested, colonization levels were approximately 10 times higher following food-borne infection (Fig. 2C). Food-borne infection also led to increased persistence of EHEC in the intestinal tract: While water-borne EHEC were gradually cleared by most fish over the course of 24 hours, food-borne EHEC persisted at high levels up to 24 hours post infection (hpi) in approximately 50% of the fish, while the other half carried lower burdens or cleared the infection over the same time span (Fig. 2D).

**Fig. 2.**
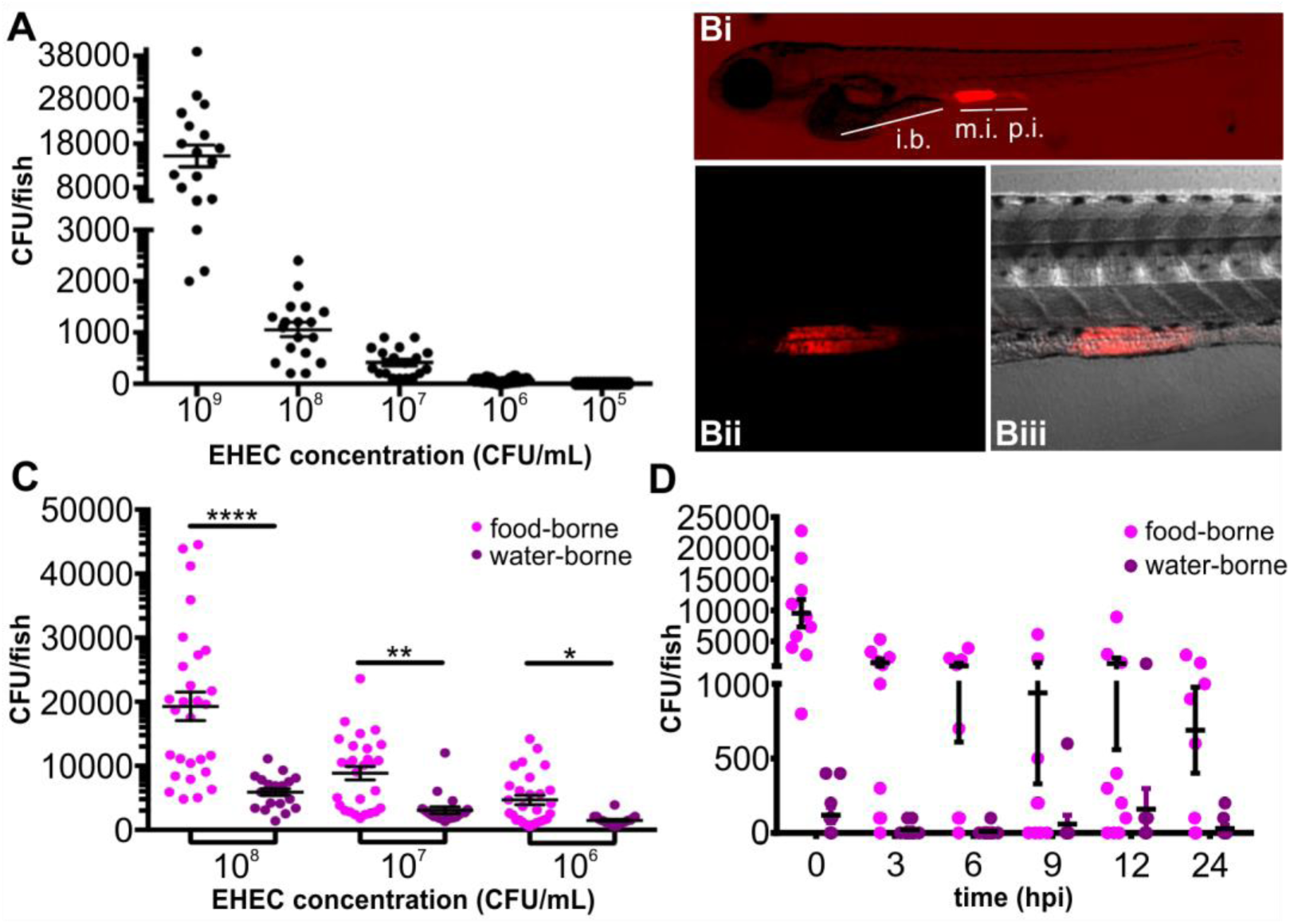
Colonization of larval zebrafish by water-borne and food-borne EHEC. (A) Water-borne EHEC infection. Zebrafish larvae were exposed to EHEC for two hours at the indicated concentrations. Following exposure, fish were washed thoroughly, homogenized, and CFU determined by dilution plating on EHEC selective agar. Individual data points, means (n=18 for each concentration) and stdev are shown. (B) Live image of zebrafish larvae exposed to water-borne EHEC:mCherry (10^8^ CFU/mL) for 2 hours. Whole embryo (i), magnified section of mid-intestinal region (EHEC:mCherry, red, (ii) and EHEC:mCherry/DIC merge (iii) are shown. i.b.: intestinal bulb, m.i.: mid-intestine, p.i.: posterior intestine. (C) Zebrafish larvae were exposed to food-borne (magenta) or water-borne (purple) EHEC for two hours, at indicated doses. Bacterial burden was determined by dilution plating on EHEC selective agar. Individual data points, means (n=28 for each condition) and stdev are shown. Statistical significance was determined using student’s t-test and defined as p< 0.05 (*), p< 0.01 (**), or p< 0.0001 (****). (D) Zebrafish larvae were exposed to 10^8^ CFU/mL of food-borne (magenta) or water-borne (purple) EHEC for two hours, transferred to fresh sterile medium, and bacterial burden was determined by dilution plating on EHEC selective agar at indicated time points. Individual data points, means (n=10 for each condition) and stdev are shown.

### EHEC triggers transient neutrophil recruitment to the mid-intestine upon colonization

EHEC infection causes release of IL-8, a potent neutrophil chemoattractant, by the intestinal mucosa (19). Humans infected with EHEC shed a high number of leukocytes in their feces (20) and tissue biopsies taken from patients infected with EHEC revealed a severe inflammatory response to EHEC, and accompanying damage of the tissue in the cecum and colon (21, 22). To test whether EHEC colonization triggers a pro-inflammatory response in zebrafish, we followed neutrophil movement in infected transgenic larvae (Tg(*mpo::egfp*)^i114^) featuring GFP-expressing neutrophils. These transgenic zebrafish express GFP under the neutrophil-specific myeloperoxidase (*mpo*) promoter (23). Imaging was limited to 12 hpi, due to technical limitations of the microscope setup. However, rapid neutrophil recruitment to the intestinal region was observed in infected, but not in uninfected control animals (Fig. 3). Neutrophils were especially concentrated in areas adjacent to the mid-intestinal infection focus and anal region, and neutrophil migration through the epithelium and into the lumen was observed (Fig. 3A). Recruitment was significantly elevated at 2 hpi, peaked around 8 hpi and declined thereafter (Fig. 3B). This was accompanied by a decline in the EHEC:mCherry fluorescence signal, suggesting that neutrophils are involved in bacterial clearance.

**Fig. 3.**
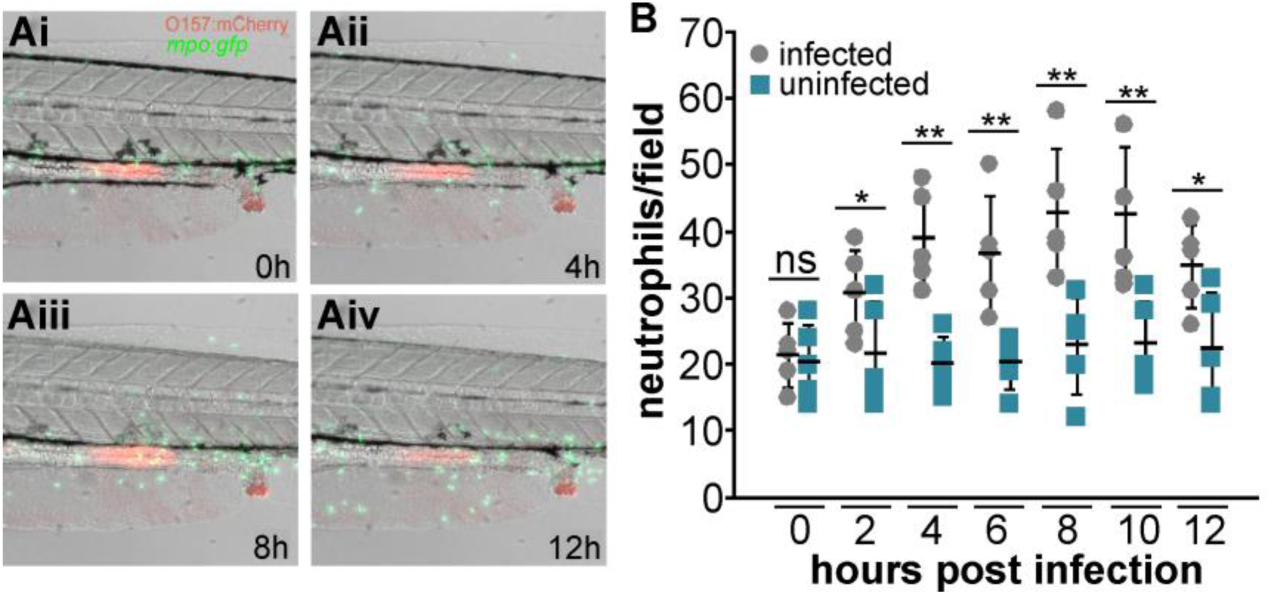
EHEC triggers transient neutrophil recruitment to the mid-intestine upon colonization. *Mpo:gfp* zebrafish larvae were exposed to 10^8^ CFU/mL food-borne EHEC:mCherry for two hours, washed, live-mounted in low-melt agarose containing tricaine, and imaged for 12 hours at 32°C. (A) Representative frames showing intestinal tract of infected fish after 0 hours (Ai), 4 (Aii), 8 (Aiii), and 12 hours (Aiv) are shown. (B) Neutrophils in the intestinal region (counts from intestinal region as shown in A) were enumerated. Individual data points from infected (grey) and uninfected fish (blue), means (n=5 per condition) and stdev are shown. Statistical significance was determined using student’s t-test and defined as p< 0.05 (*) or p< 0.01 (**). ns: not significant.

### EHEC induce the locus of enterocyte effacement at the site of colonization

The locus of enterocyte effacement (LEE) is a horizontally acquired pathogenicity island encoding for structural and regulatory components of a type three secretion system (T3SS). It is a shared feature between A/E lesion producting pathogens, including enteropathogenic *E. coli* (EPEC), *Citrobacter rodentium* and EHEC, and contributes to intestinal persistence and disease severity in animal models (5, 24). Expression of LEE encoded genes is silenced by H-NS under environmental conditions (25), and induced under physiological conditions mimicking the host intestinal tract (26). The LEE is organized in five transcriptional units, which require the LEE1 encoded masterregulator Ler (LEE encoded regulator) for induction (27). Using an EHEC:mCherry reporter strain carrying the LEE1 promoter driving GFP expresssion, we investigated if and where this virulence factor is expressed by EHEC following ingestion by zebrafish.

While EHEC LEE1:*gfp* showed low levels of fluorescence outside the host (Fig. 4A), GFP expression, and thus LEE induction, was observed in the zebrafish intestinal tract (Fig. 4B). In contrast, a control strain lacking a promoter driving GFP expression formed similar infection foci, but did not express GFP (Fig. 4C). This suggests the LEE encoded T3SS is induced during colonization of the zebrafish gastrointestinal tract.

**Figure 4.**
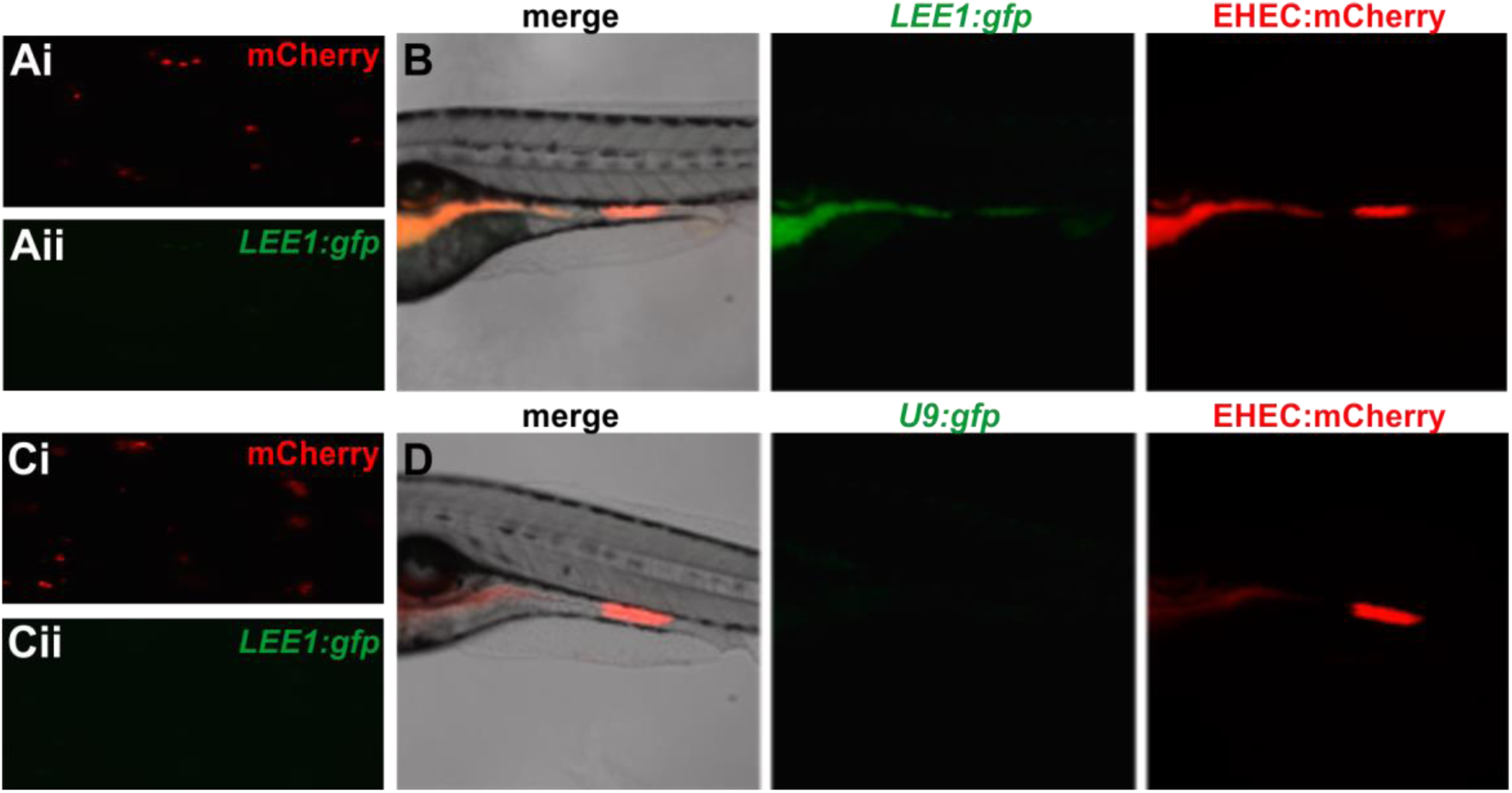
EHEC induce the locus of enterocyte effacement at the site of colonization. Micrograph of EHEC:mCherry co-transformed with *LEE1:gfp* grown in planktonic media (A) and zebrafish colonization by EHEC:mCherry / LEE1:gfp (B). EHEC:mCherry co-transformed with promoter-less control U9:gfp grown planktonically (C) and zebrafish colonization by EHEC:mCherry co-transformed with U9:gfp (D). Zebrafish were exposed to 10^9^ CFU/ml of food-borne EHEC for 2 hours.

### Fine-tuning of LEE expression is required for successful colonization

Since we observed LEE expression in the zebrafish intestinal tract, we next tested if LEE expression and EHEC intestinal colonization levels were linked, using the zebrafish model. The metabolic enzyme AdhE, a bi-functional acetaldehyde-CoA dehydrogenase and alcohol dehydrogenase, regulates virulence gene expression in EHEC O157. Deletion of *adhE* leads to strong suppression of the T3SS (28). The *adhE* mutant is significantly less virulent than its parental strain, TUV 93-0, and showed approximately ten-fold lower colonization levels in an infant rabbit infection model (28). These observations from the infant rabbit model were recapitulated in our zebrafish model: Following a two hour exposure to 10^8^ CFU/mL of food-borne EHEC, LEE1:*gfp* was significantly induced in the TUV 93-0 wild type strain colonizing the instestinal tract, compared to plantonic growth (Fig. 5A, B). In contrast, reduced LEE1:*gfp* induction was observed in the *adhE* mutant, in both planktonic culture or during host colonization, as analyzed by imaging (Fig. 5C, D) and ratiometric analysis across several colonized fish (Fig. 5E). The bacterial burden was approximately ten-fold reduced for the *adhE* mutant compared to the TUV 93-0 wild type (Fig. 5F), although both growth (Fig. 5G) and persistence within *P. caudatum* (Fig. 5H) were unaltered.

**Figure 5.**
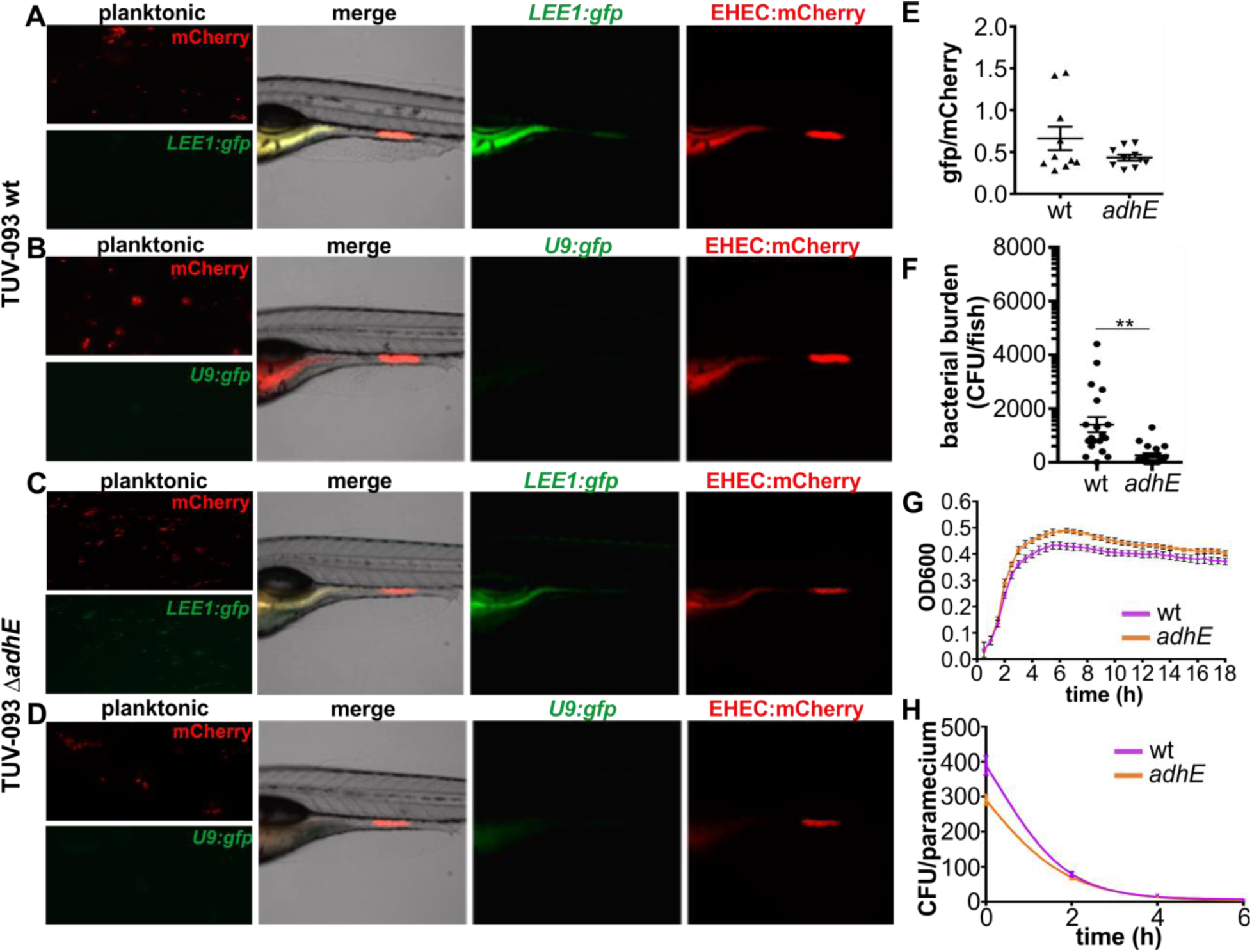
LEE expression and colonization of EHEC Δ*adhE*. (A) EHEC TUV 93-0 wild type transformed with mCherry and *LEE1:gfp* grown planktonically and in zebrafish. (B) EHEC TUV 93-0 wild type transformed with mCherry and promoter-less *U9:gfp* grown planktonically and in zebrafish. (C) EHEC TUV 93-0 isogenic Δ*adhE* mutant transformed with mCherry and *LEE1:gfp* grown planktonically and in zebrafish. (D) EHEC TUV 93-0 isogenic Δ*adhE* mutant transformed with mCherry and promoter-less *U9:gfp* grown planktonically and in zebrafish. (E) Ratiometric image analysis comparing levels of *LEE1:gfp* expression normalized to mCherry signal for EHEC TUV 93-0 wild type and Δ*adhE* mutant (n=9 fish/condition). (F) Bacterial burden was determined by dilution plating on EHEC selective agar. Individual data points, means (n=15 for each condition) and stdev are shown. Statistical significance was determined using student’s t-test; p< 0.01 (**). (G) Growth curves of TUV 93-0 wild type and Δ*adhE* mutant. (H) Degradation profiles for TUV 93-0 wild type and Δ*adhE* mutant in *P. caudatum* as determined by lysis and dilution plating. Regression analysis showed there is no difference between the slopes (p=0.2909).

Next, we set out to test the effect of enhanced T3SS expression on colonization levels and persistence of EHEC in the zebrafish gut, using the *LEE1:gfp* reporter. The protein TolA is a component of the trans-envelope Tol system required for cell wall remodeling in *E. coli* (29). Loss of *tolA* in EHEC leads to activation of the Rcs phosphorelay, and constitutive activation of LEE encoded virulence genes (30). Despite a higher level of T3SS expression in Δ*tolA*, the strain’s adherence to intestinal epithelial cells and its virulence in a *Galleria mellonella* infection model are significantly reduced (30). We constructed a non-polar *tolA* mutant in the Sakai background, and compared its LEE induction levels and ability to colonize the zebrafish intestine to those of the wild type Sakai strain following two hours of infection. *LEE1:gfp* transcriptional activity was significantly higher in planktonically grown *tolA* compared to wild type bacteria, as previously described (30). This also held true for *tolA* within the zebrafish intestinal tract: Although *LEE1:gfp* activity relative to the constitutively active mCherry was enhanced for *tolA* compared to wild type bacteria within zebrafish, the bacterial burden of the *tolA* mutant was significantly reduced compared to the wild type (Fig. 4 and Fig. 6A-F). This was not due to altered growth or degradation within the *P. caudatum* vector (Fig. 6G, H). Overall, these data suggest that the tight regulation of T3SS expression is essential for within-host fitness.

**Figure 6.**
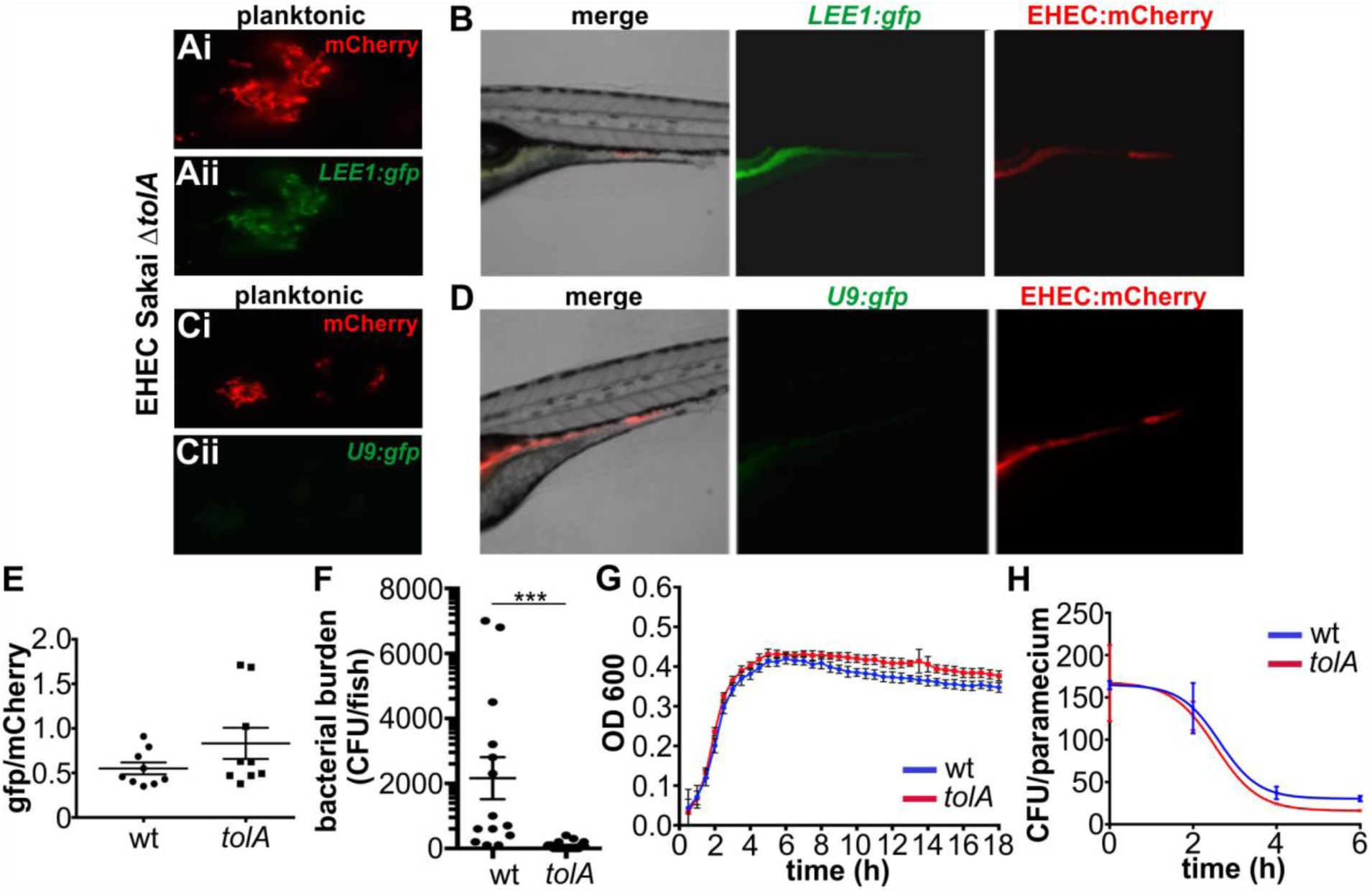
LEE expression and colonization of EHEC Δ*tolA*. EHEC Sakai isogenic Δ*tolA* mutant transformed with mCherry and *LEE1:gfp* grown planktonically (A) and in zebrafish (B). EHEC Sakai isogenic Δ*tolA* mutant transformed with mCherry and promoter-less *U9:gfp* grown planktonically (C) and in zebrafish (D). (E) Ratiometric image analysis comparing levels of *LEE1:gfp* expression normalized to mCherry signal for EHEC Sakai wild type and Δ*tolA* mutant (n=10 fish/condition). (F) Bacterial burden was determined by dilution plating on EHEC selective agar. Individual data points, means (n=15 for each condition) and stdev are shown. Statistical significance was determined using student’s t-test; p< 0.001 (***). (G) Growth curves of Sakai wild type and Δ*tolA* mutant. (H) Degradation profiles for Sakai wild type and Δ*tolA* mutant in *P. caudatum* as determined by lysis and dilution plating. Regression analysis showed there is no difference between the slopes (p=0.7016).

### EHEC ingestion causes strain-specific mortality in zebrafish

Having established that T3SS expression is specifically induced in the zebrafish host and is necessary for efficient intestinal colonization, we asked if EHEC ingestion would cause mortality in zebrafish. We infected zebrafish at 4 dpf with a dose of 10^9^ CFU/ml of food-borne EHEC or left them uninfected (control). Fish were assessed for vital signs (movement, heart beat and circulation) daily for a total of six days post infection (dpi), and survival was analysed using the Kaplan-Meier estimator (Fig. 7). Despite showing similar colonization levels (Fig. 5 and 6), Sakai and TUV 93-0 wild type strains displayed significantly different pathogenicity, with a mean survival of 76% for Sakai and 56% for TUV 93-0 at the experimental endpoint. The attenuation of the *tolA* and *adhE* mutants observed in terms of intestinal colonization (Fig. 4-6) was also reflected by a reduction in pathogenicity. The *tolA* mutant was non-pathogenic in the zebrafish model, with 100% host survival and no observable morbidity up until day 6 post infection. The *adhE* mutant was similarly attenuated, with a mean host survival of 87% at day 6 post infection (Fig. 7).

**Figure 7.**
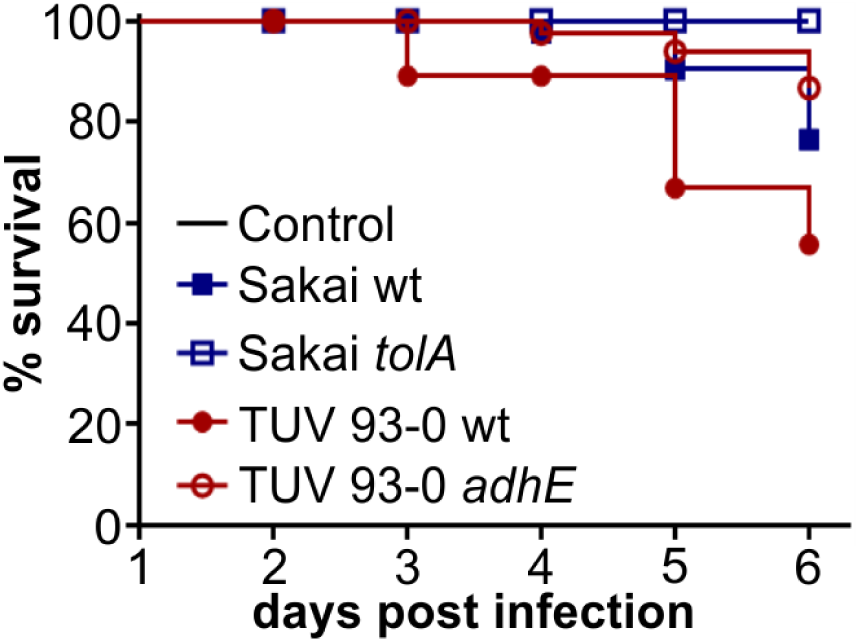
EHEC ingestion causes strain-specific mortality in zebrafish. Zebrafish at 4dpf were infected with 10^9^ CFU/ml of EHEC (Sakai wt - blue full squares, Sakai *tolA* – empty squares, TUV 93-0 wt – red filled circles, or TUV *adhE* – red empty circles) or left uninfected (black line, runs below *tolA* data), and incubated at 32 °C for up to 6 dpi. Fish were assessed for vital signs (presence of either movement, heart beat or circulation) every 24 hours and percent survival plotted using Kaplan-Meier analysis (n=15 per condition).

### Zebrafish as a model to study fecal shedding and transmission

While fecal shedding of EHEC is often used as a proxy for bacterial burden in rodents, fecal-oral transmission is rarely studied in these models. The zebrafish model allows simultanous analysis of fecal shedding and fecal-oral transmission from infected fish to naïve recipients in one experiment. AB fish infected with food-borne EHEC for two hours were transferred into fresh media together with a naïve recipient (Fig. 8A). Tg(*mpo:gfp*) fish were used as recipients to be able to visually distinguish donor and recipient (green fluorescent). EHEC was continutally shed from donor fish following transfer into fresh media, and levels of shed bacteria increased steadily up until 24 hours post transfer (Fig. 8B). Onward transmission to naïve fish was first observed between 12-24 hours post transfer (Fig. 8C), at which point EHEC:mCherry could be visualized in the foregut and mid-intestine of recipient embryos (Fig. 8D). These data demonstrate that the dynamics of fecal shedding and fecal-oral transmission to naïve hosts can be studied using the zebrafish model.

**Figure 8.**
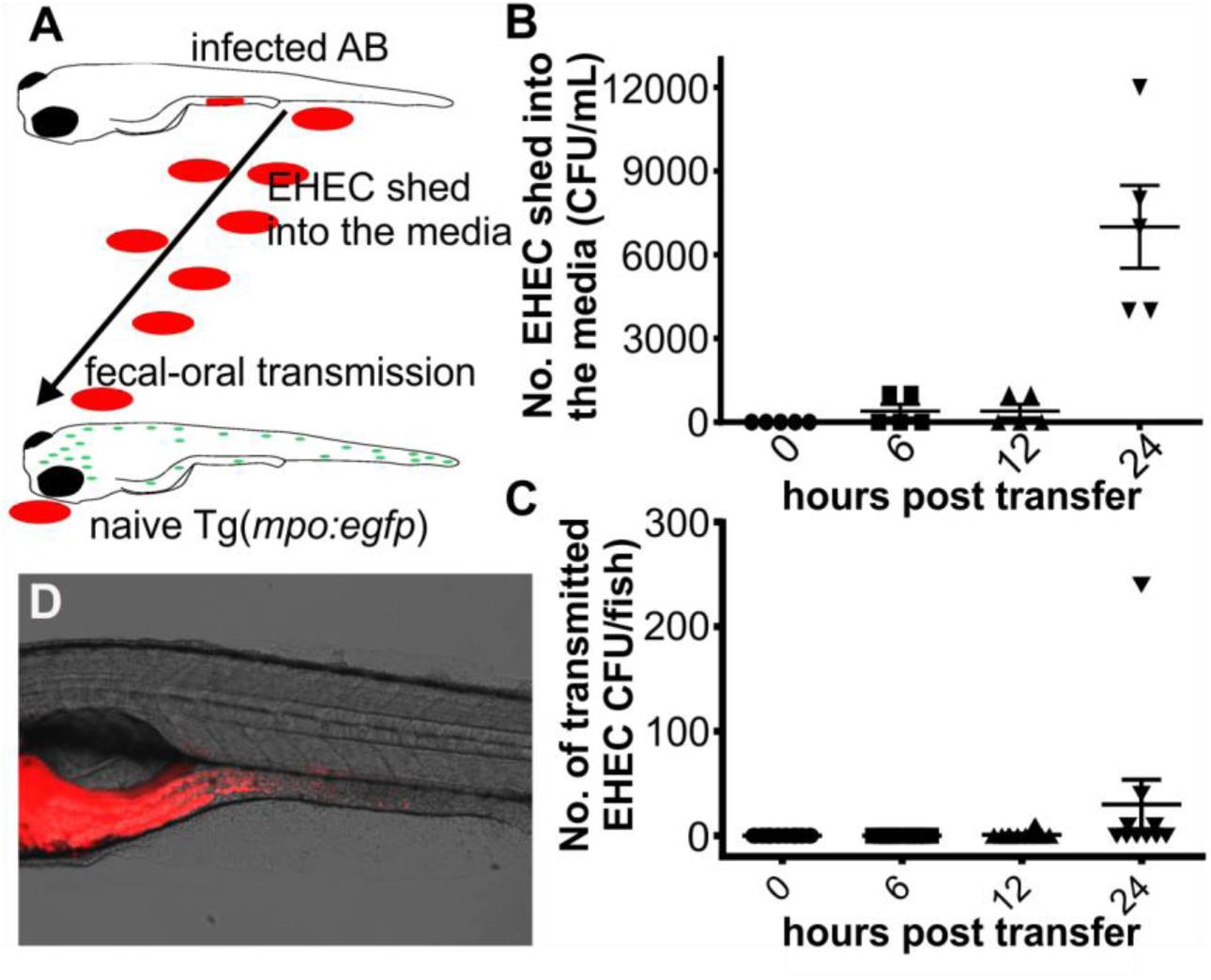
Zebrafish as a model to study fecal shedding and transmission. (A) Scheme depicting experimental setup of shedding and transmission experiment. Infected AB embryos were transferred into fresh media housing naïve Tg(*mpo:egfp*) embryos (both 4 dpf). Post transfer, shedding of EHEC into the media and transfer to recipient animals were monitored over time. (B) EHEC shed into the media were enumerated immediately, 6, 12 or 24 hours post transfer of infected AB embryos into fresh media. Shown are individual data points, means and stdev (n=5/time point). (C) Bacterial burden of Tg(*mpo:egfp*) recipient embryos was quantified immediately, 6, 12 and 24 hours post transfer of infected AB (donor) embryos. Shown are individual data points, means and stdev (n=10 embryos/time point). (D) Example of EHEC:mCherry localization in recipient embryos.

### The zebrafish microbiota provides a barrier to EHEC colonization

To this point, our experiments were done in gnotobiotic zebrafish, as the endogenous microbiota often proves a significant barrier to colonization (31, 32). Yet, the interactions between ingested pathogens and the endogenous microbiota, and the impact of the microbiota composition on the fate of infection, are of interest, particularly in the case of EHEC (33-35), and it would be desirable to be able to address these questions in the zebrafish infection model.

It has been reported that zebrafish acquire a microbiota which rapidly diversifies during early development (18, 36, 37). To test whether EHEC infection can be studied in the zebrafish model against the backdrop of the endogenous microbiota, we compared levels of EHEC in gnotobiotic fish and conventionalized fish, which were transferred into a mixture of E3 and tank water following hatching. Compared to gnotobiotic fish, initial colonization with EHEC was significantly decreased, but not entirely eliminated, in conventionalized fish (Fig. 9A).

Surprisingly, colonization levels in conventionalized fish were very consistent, even though the composition as well as levels of the colonizing microbiota differed considerably between individual animals (Fig. 9B). In conventialized fish, the bacterial burden expanded with increased incubation time, and could easily be visualized from 16 hpi (Fig. 9C).

**Fig. 9.**
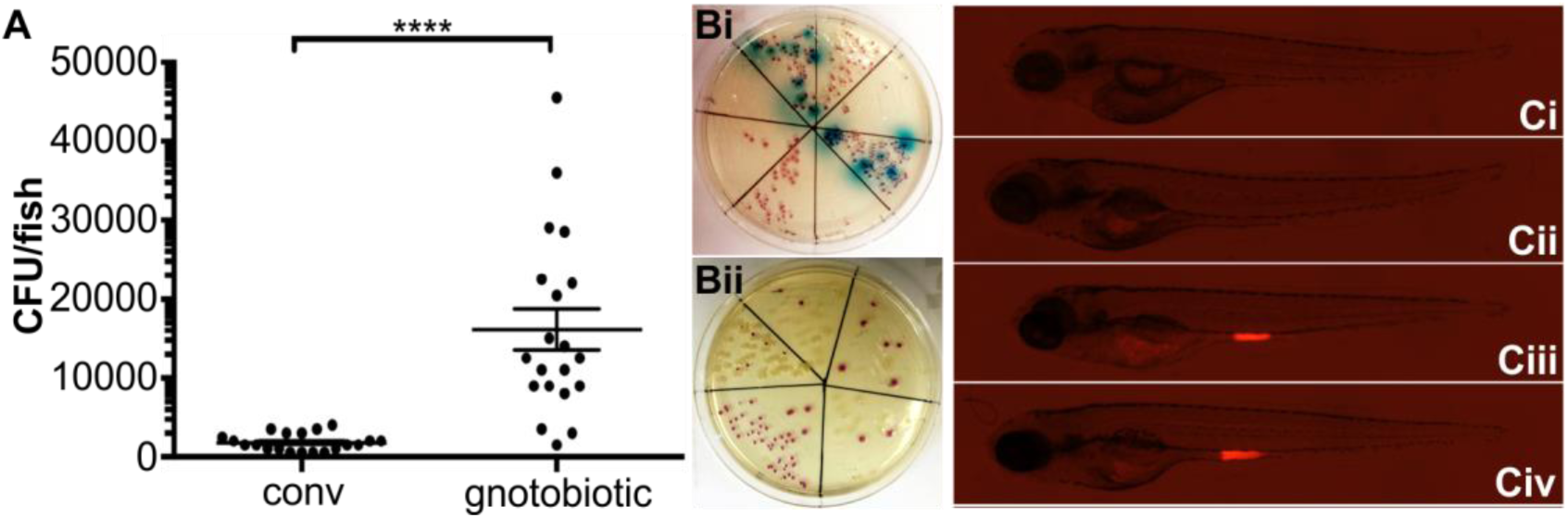
The zebrafish microbiota provides a barrier to *E. coli* colonization. (A) Following exposure of conventionalized or gnotobiotic zebrafish larvae to food-borne EHEC (10^8^ CFU/mL) for two hours, bacterial burden was determined by dilution plating on EHEC selective agar. Individual data points, means (n=20 for each condition) and stdev are shown. Statistical significance was determined using student’s t-test, and defined as p< 0.0001 (****). (B) Examples of larval homogenates derived from conventionalized fish following EHEC infection, plated on EHEC selective CHROMagar. Mauve, blue and white colonies correspond to EHEC, other coliforms, and *Proteus sp*, respectively. (C) Colonization of conventionalized zebrafish with EHEC:mCherry following 2 (Ci), 8 (Cii), 16 (Ciii) and 24 hours (Civ) of infection.

## DISCUSSION

Here, we establish zebrafish larvae as a new vertebrate model for EHEC infection. Maintenance of zebrafish is inexpensive and their propagation and development is quick compared to other vertebrate animals, making them an attractive model organism for infection biology. Zebrafish have been used extensively to study bacterial infections, but to date few gastrointestinal infection models have been described and our study provides, to our knowledge, the first extensive description of a food-borne infection model in zebrafish. *P. caudatum* is an ideal vector for food-borne infections: It is commonly used as a food-source for young zebrafish, and interactions between *E. coli* and *Paramecia* have been characterized previously (38). In agreement with earlier studies, we found no detrimental effect of EHEC-associated virulence factors on *P. caudatum* proliferation. *E. coli* is taken up into food vacuoles within seconds to minutes, depending on the density of suspended particles (39). Vacuoles gradually acidify from an initial pH of 8 to reach a pH of close to 1 (39), and EHEC have a half-life of approximately 150 minutes under those conditions. This bacterial passage through an acidifying compartment and subsequent release into the zebrafish foregut upon ingestion of paramecia mimicks passage through the human GI tract and acidification in the stomach.

Upon ingestion, EHEC rapidly colonize the mid-intestine, and although bacteria were initially observed both in fore-gut and mid-intestinal tract, EHEC shows a distinct preference for colonizing the mid-intestine, and the site of infection displays sharp boundaries, with temporary colonization of the fore-gut during early infection, and no colonization of the posterior intestine (Fig. 2, 4). In contrast, colonization of infant rabbit is not as localized, with similar bacterial burdens found in ileum, cecum and colon even at later time points (5).

Human EHEC infection is known to cause a strong pro-inflammatory response, and neutrophil infiltration of the lamina propria and transmigration through the intestinal epithelium into the gut lumen has been described in monkey, piglet, and rodent models (5, 40, 41). This is a response of increased IL-8 production by the intestinal mucosae, although it has been a point of contention whether Stx or TLR5 recognition of H7 flagellin was the major factor inducing IL-8 secretion. I*n vitro* and *ex vivo* studies using flagellar mutants demonstrated that IL-8 secretion is driven by exposure of epithelial cells to flagellar antigen, and to some extent, TLR4-driven responses to LPS (42, 43). In our model, Stx negative EHEC is still capable of mounting a strong neutrophilic inflammation, which supports these data. While neutrophil depletion has been shown to increase the bacterial burden of *Citrobacter rodentium* in mouse (44) and targeting leukocyte adhesion factor with antibody reduces disease symptoms in rabbits (3), no direct link between neutrophil recruitment and bacterial clearance has been established for EHEC. The zebrafish immune system displays many similarities to that of mammals, with counterparts for most human immune cell types (45). The zebrafish innate immune system starts to develop as early as 24 hpf with primitive macrophages followed by neutrophils at 32-48 hpf. The development of the adaptive immune system lags behind, taking another 4 weeks to fully develop (46). This feature provides an opportunity to exclusively observe the innate immune reaction to an EHEC infection in our experimental system. Real-time imaging of EHEC infected zebrafish allowed us to simultaneously follow and establish a temporal link between neutrophilic inflammation and EHEC persistence in the intestine. We observed a significant increase in neutrophil recruitment to the infection site as early as 2 hpi, with a peak around 8 hpi. After that, the response gradually diminished until the experimental endpoint (12 hpi). This coincided with bacterial burden, which was significantly decreased 3 hpi, and gradually diminished thereafter. However, a low level of persistence was still observed at the experimental endpoint, 24 hpi. Although only circumstantial, the timing of these two events suggests bacterial clearance may, at least in part, be mediated by neutrophils.

EHEC infection in infant rabbits is self-limiting, with a peak in bacterial burden approximately 7 dpi and a decline in inflammation and bacterial burden thereafter (5). We observe a similar, albeit accelerated pattern in zebrafish, with peak inflammation approximately 8 hpi. Bacterial burden declines over time, and interestingly, there are two distinct colonization patterns:

Approximately half the animals clear the infection, while the other half display low but persistent colonization until the experimental endpoint, without signs of clearance (Fig. 2D). Although current licence requirements prevented us from doing so, future work will focus on following infections for longer, to establish if EHEC is eventually fully cleared from these animals.

Following attachment to the host epithelia, EHEC regulates the cordinated expression of a range of bacterial effectors through the activity of LEE 1-5 encoded virulence factors. The LEE1 encoded masterregulator of the LEE pathogenicity isleand, Ler, has previously been shown in *in vitro* models to be activated early in infection and is regulated in part through bacterial attachment and sensing of fluid shear (47). The activity of the five LEE operons is also temporal, with *in vitro* models demonstrating that by 3 hpi LEE1 activity is down regulated, while transcription of LEE3 and LEE5 encoded genes is upregulated (48). Through the use of EHEC strains expressing a LEE1:*gfp* reporter we were able to utilise the zebrafish model to visualise LEE1 activation *in vivo* during the early stages of infection, and correlate its expression with the EHEC colonization pattern. Our results demonstrate the importance of tight control of EHEC virulence gene regulation for successful colonization *in vivo*. Deletion of *adhE* which post-transcriptionally affects the regulation of virulence genes through Hfq, suppresses production of the LEE-encoded T3SS and decreases colonization in a rabbit model (28). Decreased colonization was also recapitulated in the zebrafish model, with both LEE1 induction and bacterial burden decreased in an *adhE* mutant compared to the TUV 93-0 wild type strain (Fig. 5). Our data further demostrate that temporal regulation of LEE activation is important to success of colonization: Deletion of *tolA* causes constitutive activation of the Rcs phosphorelay and expression of LEE encoded virulence genes outside of the host environment (30). Despite overexpression of T3SS, *tolA* mutants display decreased attachment to intestinal epithelial cells and a reduced virulence in *Galleria mellonella* infection models. Our model recapitulated these findings, in that despite higher levels of LEE1 expression, both in planktonic culture and *in vivo*, the bacterial burden was significantly reduced compared to the Sakai wild type strain (Fig. 6). Both *adhE* and *tolA* strains cause significantly reduced morbidity in the zebrafish model, compared to wild type strains (Fig. 7). These findings demonstrate that the zebrafish system faithfully reflects the contribution for T3SS, a key virulence factor, and its regulation to bacterial colonization and persistence. Our data also underpin that induction of virulence factors has to be carefully timed, and fine-tuned in response to the host environment with deregulated induction of virulence genes being just as detrimental to a pathogen’s fitness as loss of virulence factors.

In summary, the zebrafish represents a powerful model to study EHEC infection. The optical clarity of zebrafish larvae and the availablity of transgenic fish lines, coupled to the use of bacterial reporter strains, enables simultaneous visualisation of bacterial physiology and the molecular dynamics of host-pathogen interactions in real time within a living organism. This model will therfore provide oportunities to study the molecular aspects governing EHEC colonization and pathogenesis *in vivo,* at a single cell level and with refined temporal resolution and sampling size.

## MATERIALS AND METHODS

### Bacterial strains and growth conditions

Bacterial wild type strains used in this study were EHEC O157:H7 strain 813, a derivative of *Eschericha coli* O157:H7 Sakai (a gift from S. Busby) and *Eschericha coli* O157:H7 TUV 93-0 (a gift from A. Roe), both Stx negative. The Δ*tolA* mutant was a derivative of strain 813 and generated by gene-doctoring (49). Mutant Δ*adhE* (28, 50), was derived from TUV 93-0 (51), and received as a gift from A. Roe (Univ. of Glasgow). For monitoring localization *in vivo*, strains were transformed with plasmid pDP151-mCherry (ampR). To visualize induction of the LEE1-encoded regulator Ler, strains were transformed with plasmid LEE1:*gfp* or U9:*gfp* (promoter-less negative control), both tetR (47). **S**trains were grown in LB broth supplemented with 100 μg/ml ampicillin or 34 μg/ml tetracyclin, where appropriate, at 37 °C with gentle shaking.

### Fish maintenance and breeding

Zebrafish (*Danio rerio*) strains used in this study were AB wildtype fish and transgenic fish of the Tg(*mpo::egfp*)^i114^ line that produce green fluorescent protein (GFP) in neutrophils (23). Adult fish were kept in a recirculating tank system at the University of Birmingham Aquatic Facility, or the UTHealth Center for Laboratory Animal Medicine and Care at a 14/10 hours light/dark cycle at a pH of 7.5 and 26 °C. Zebrafish care, breeding and experiments were performed in accordance with the Animal Scientific Procedures Act 1986, under Home Office Project License 40/3681 (University of Birmingham) and Animal Welfare Committee protocol AWC-16-0127. Eggs were obtained from natural spawning between adult fish, which were set up in groups of seven (4♀/3♂) in separate breeding tanks. After collection of eggs, larvae were kept in a diurnal incubator under a 14h-10h light-dark cycle with the temperature maintained at 28.5 °C. Embryos were raised in petri dishes containing E3 medium (5 mM NaCl, 0.17 mM KCl, 0.33 mM CaCl_2_, 0.33 mM MgSO_4_) supplemented with 0.3 μg/ml of methylene blue. From 24 hpf, 0.003 % 1-phenyl-2-thiourea (PTU) was added to prevent melanin synthesis. During infections, larvae were maintained at 32 °C. All zebrafish care and husbandry procedures were performed as described previously (52).

### Preparation and maintenance of gnotobiotic and conventionalized fish

Gnotobiotic embryos were produced by step-wise bleaching of eggs with a 0.0045 % sodium hypochlorite solution in E3 at 12 hpf, as previously described (52). Since the sodium hypochlorite solution leads to a hardening of the chorion, embryos were freed manually by dechorination. After bleaching, eggs were transferred and maintained in sterile E3 medium. To enable colonization of embryos by a conventional gut microbiota, embryos were transferred to untreated tank water directly obtained from the aquatic system used for maintenance of adult zebrafish following dechorination.

### Maintenance of *Paramecium caudatum*

Paramecia were cultured at 22 °C in 5 cm petri dishes with E3 medium containing *E. coli* MG1655 as a food source. To maintain the culture, 0.5 ml of an existing paramecium culture were passaged into 9.5 mL of fresh E3 medium containing 10^8^ CFU/mL of *E. coli* MG1655.

### Uptake and clearance of *Escherichia coli* O157:H7 by P*. caudatum*

The uptake of fluorescent EHEC by paramecia into vacuoles was observed by live imaging and on fixed samples under a fluorescent microscope. The acidification of the food vacuoles and subsequent degradation/clearance of bacteria was followed by an additional staining with lysotracker red. To determine EHEC viability within *P. caudatum*, samples were removed from *P. caudatum* cultures at indicated time points, lysed with 1% Triton-X100 in PBS and homogenized, and serial dilutions in PBS plated on CHROMagar™ O157 plates for enumeration of CFUs.

### Food- and water-borne fish infections

For infection experiments bacteria were harvested by centrifugation at 6000 g for 2 min and adjusted to an OD_600_ of 1.0 (≅ concentration of 2x10^8^ bacteria/ml). *P. caudatum* were quantified using a hemocytometer and added to the suspension to give a concentration of 10^4^ paramecia/mL, and incubated for 2 h at 32 °C. Following pre-incubation, paramecia were washed and added to zebrafish larvae (4 dpf) housed in 6-well plates, to give the indicated bacterial concentrations. For water-borne infections, EHEC concentrations as indicated were directly prepared in E3 and added to zebrafish larvae (2ml/10 zebrafish larvae/6-well). Following infections, zebrafish larvae were either anaesthetized by adding tricaine (final conc. 160 μg/mL) to 20 mL E3 and 2 mL of a 0.1 M sodium-bicarbonate solution to buffer the medium, or euthanized by adding 1.6 mg/mL tricaine to the buffered medium. For microscopy, larvae were transferred to a 1.5 mL microcentrifuge tube, washed in PBS, fixed by adding 1 mL of a 4 % para-formaldehyde solution in PBS, and stored at 4 °C in the dark until used.

### Imaging of infected fish

For live imaging, infected anaesthetized larvae were positioned in 96-well glass-bottom plates and covered and immobilized with 1 % low-melting-point agarose solution. 200 μl E3 containing 160 μg/mL tricaine was added to cover the immobilized larvae. Live imaging was performed at 32 °C and 80 % humidity. A Zeiss Axio Observer inverse microscope equipped with a 10x objective was used for acquisition of 2 fluorescent channels and 1 differential interference contrast (DIC) channel. The 4D images produced by the time-lapse acquisitions were processed, clipped, examined and interpreted using the Zen 2 software (Zeiss). Maximum intensity projection was used to project developed Z-stacks and files were exported in tiff format for images or mov format for QuickTime movies. Final figures were assembled with CorelDrawX5.

### Determination of bacterial burden in infected fish

After euthanasia larvae were transferred to individual microcentrifuge tubes and disintegrated by repeated pipetting and vortexing in 200 μl PBS supplemented with 1% Triton X-100. Serial dilutions of bacterial suspension were plated onto selective CHROMagar™ O157 plates and plates were incubated at 37 °C for 16 hours. Colonies identified as EHEC (mauve appearance on selection plate) were quantified for analysis.

## ACKNOWLEDGMENTS

We thank A. Roe for sharing strain TUV 93-0 and its derivatives. We thank S. Renshaw, S. Johnston and R. Wheeler for sharing Tg(*mpo:gfp*) eggs. We thank members of the Krachler and Voelz labs for critical reading and comments on the manuscript.

---

**Fig. S1.**
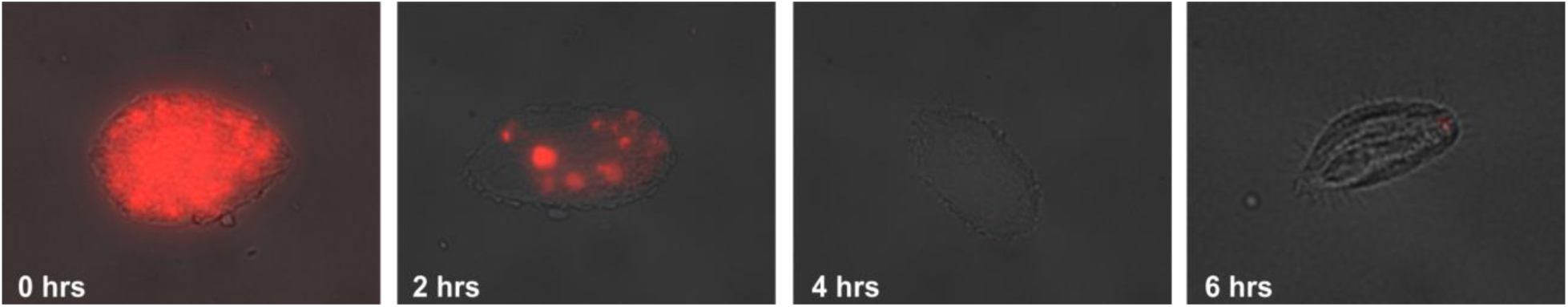
Degradation of internalized EHEC by *P. caudatum.* At specified time points, paramaecia incubated with EHEC Sakai:mCherry were washed, fixed in 4% PFA, mounted in anti-fade Gold and imaged using the Zeiss Axio Observer.

**Figure. S2.**
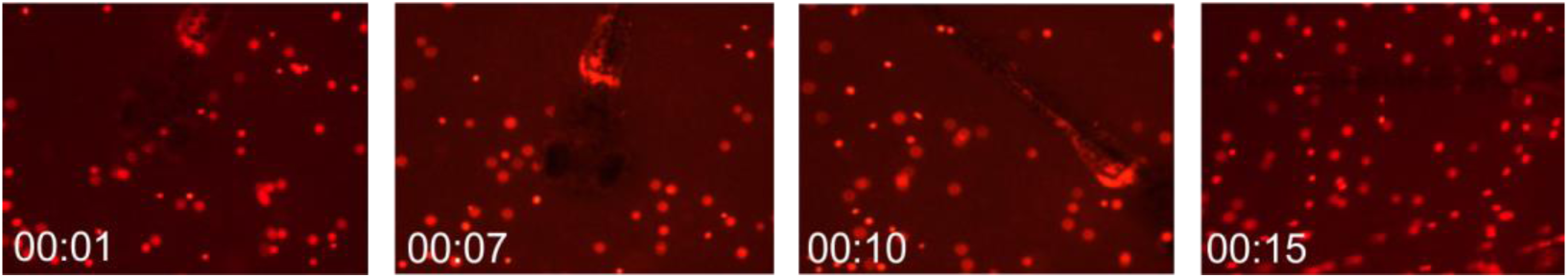
Stills from Movie S1, showing a zebrafish larva at 4 dpf preying on *P. caudatum* loaded with EHEC Sakai:mCherry.

## REFERENCES

1. Donnenberg MS, Tacket CO, James SP, Losonsky G, Nataro JP, Wasserman SS, Kaper JB, Levine MM. 1993. Role of the eaeA gene in experimental enteropathogenic Escherichia coli infection. J Clin Invest 92:1412–7.

2. Ritchie JM. 2014. Animal Models of Enterohemorrhagic Escherichia coli Infection. Microbiol Spectr 2:EHEC-0022-2013.

3. Elliott E, Li Z, Bell C, Stiel D, Buret A, Wallace J, Brzuszczak I, O'Loughlin E. 1994. Modulation of host response to Escherichia coli o157:H7 infection by anti-CD18 antibody in rabbits. Gastroenterology 106:1554–61.

4. Ogawa M, Shimizu K, Nomoto K, Takahashi M, Watanuki M, Tanaka R, Tanaka T, Hamabata T, Yamasaki S, Takeda Y. 2001. Protective effect of Lactobacillus casei strain Shirota on Shiga toxin-producing Escherichia coli O157:H7 infection in infant rabbits. Infect Immun 69:1101–8.

5. Ritchie JM, Thorpe CM, Rogers AB, Waldor MK. 2003. Critical roles for stx2, eae, and tir in enterohemorrhagic Escherichia coli-induced diarrhea and intestinal inflammation in infant rabbits. Infect Immun 71:7129–39.

6. Wadolkowski EA, Burris JA, O'Brien AD. 1990. Mouse model for colonization and disease caused by enterohemorrhagic Escherichia coli O157:H7. Infect Immun 58:2438–45.

7. Pagan AJ, Yang CT, Cameron J, Swaim LE, Ellett F, Lieschke GJ, Ramakrishnan L. 2015. Myeloid Growth Factors Promote Resistance to Mycobacterial Infection by Curtailing Granuloma Necrosis through Macrophage Replenishment. Cell Host Microbe 18:15–26.

8. Mostowy S, Boucontet L, Mazon Moya MJ, Sirianni A, Boudinot P, Hollinshead M, Cossart P, Herbomel P, Levraud JP, Colucci-Guyon E. 2013. The zebrafish as a new model for the in vivo study of Shigella flexneri interaction with phagocytes and bacterial autophagy. PLoS Pathog 9:e1003588.

9. Lopez-Munoz A, Liarte S, Gomez-Gonzalez NE, Cabas I, Meseguer J, Garcia-Ayala A, Mulero V. 2015. Estrogen receptor 2b deficiency impairs the antiviral response of zebrafish. Dev Comp Immunol 53:55–62.

10. Bojarczuk A, Miller KA, Hotham R, Lewis A, Ogryzko NV, Kamuyango AA, Frost H, Gibson RH, Stillman E, May RC, Renshaw SA, Johnston SA. 2016. Cryptococcus neoformans Intracellular Proliferation and Capsule Size Determines Early Macrophage Control of Infection. Sci Rep 6:21489.

11. Rolig AS, Parthasarathy R, Burns AR, Bohannan BJ, Guillemin K. 2015. Individual Members of the Microbiota Disproportionately Modulate Host Innate Immune Responses. Cell Host Microbe 18:613–20.

12. Rendueles O, Ferrieres L, Fretaud M, Begaud E, Herbomel P, Levraud JP, Ghigo JM. 2012. A new zebrafish model of oro-intestinal pathogen colonization reveals a key role for adhesion in protection by probiotic bacteria. PLoS Pathog 8:e1002815.

13. Veneman WJ, Stockhammer OW, de Boer L, Zaat SA, Meijer AH, Spaink HP. 2013. A zebrafish high throughput screening system used for Staphylococcus epidermidis infection marker discovery. BMC Genomics 14:255.

14. Miller JD, Neely MN. 2005. Large-scale screen highlights the importance of capsule for virulence in the zoonotic pathogen Streptococcus iniae. Infect Immun 73:921–34.

15. Meijer AH, van der Vaart M, Spaink HP. 2014. Real-time imaging and genetic dissection of host-microbe interactions in zebrafish. Cell Microbiol 16:39–49.

16. Watanabe K, Nakao R, Fujishima M, Tachibana M, Shimizu T, Watarai M. 2016. Ciliate Paramecium is a natural reservoir of Legionella pneumophila. Sci Rep 6:24322.

17. Peterson TS, Ferguson JA, Watral VG, Mutoji KN, Ennis DG, Kent ML. 2013. Paramecium caudatum enhances transmission and infectivity of Mycobacterium marinum and M. chelonae in zebrafish Danio rerio. Dis Aquat Organ 106:229–39.

18. Rawls JF, Samuel BS, Gordon JI. 2004. Gnotobiotic zebrafish reveal evolutionarily conserved responses to the gut microbiota. Proc Natl Acad Sci U S A 101:4596–601.

19. Dahan S, Busuttil V, Imbert V, Peyron JF, Rampal P, Czerucka D. 2002. Enterohemorrhagic Escherichia coli infection induces interleukin-8 production via activation of mitogen-activated protein kinases and the transcription factors NF-kappaB and AP-1 in T84 cells. Infect Immun 70:2304–10.

20. Slutsker L, Ries AA, Greene KD, Wells JG, Hutwagner L, Griffin PM. 1997. Escherichia coli O157:H7 diarrhea in the United States: clinical and epidemiologic features. Ann Intern Med 126:505–13.

21. Griffin PM, Olmstead LC, Petras RE. 1990. Escherichia coli O157:H7-associated colitis. A clinical and histological study of 11 cases. Gastroenterology 99:142–9.

22. Shigeno T, Akamatsu T, Fujimori K, Nakatsuji Y, Nagata A. 2002. The clinical significance of colonoscopy in hemorrhagic colitis due to enterohemorrhagic Escherichia coli O157:H7 infection. Endoscopy 34:311–4.

23. Renshaw SA, Loynes CA, Trushell DM, Elworthy S, Ingham PW, Whyte MK. 2006. A transgenic zebrafish model of neutrophilic inflammation. Blood 108:3976–8.

24. Ritchie JM, Waldor MK. 2005. The locus of enterocyte effacement-encoded effector proteins all promote enterohemorrhagic Escherichia coli pathogenicity in infant rabbits. Infect Immun 73:1466–74.

25. Sanchez-SanMartin C, Bustamante VH, Calva E, Puente JL. 2001. Transcriptional regulation of the orf19 gene and the tir-cesT-eae operon of enteropathogenic Escherichia coli. J Bacteriol 183:2823–33.

26. Kenny B, Abe A, Stein M, Finlay BB. 1997. Enteropathogenic Escherichia coli protein secretion is induced in response to conditions similar to those in the gastrointestinal tract. Infect Immun 65:2606–12.

27. Mellies JL, Elliott SJ, Sperandio V, Donnenberg MS, Kaper JB. 1999. The Per regulon of enteropathogenic Escherichia coli: identification of a regulatory cascade and a novel transcriptional activator, the locus of enterocyte effacement (LEE)-encoded regulator (Ler). Mol Microbiol 33:296–306.

28. Beckham KS, Connolly JP, Ritchie JM, Wang D, Gawthorne JA, Tahoun A, Gally DL, Burgess K, Burchmore RJ, Smith BO, Beatson SA, Byron O, Wolfe AJ, Douce GR, Roe AJ. 2014. The metabolic enzyme AdhE controls the virulence of Escherichia coli O157:H7. Mol Microbiol 93:199–211.

29. Gray AN, Egan AJ, Van't Veer IL, Verheul J, Colavin A, Koumoutsi A, Biboy J, Altelaar AF, Damen MJ, Huang KC, Simorre JP, Breukink E, den Blaauwen T, Typas A, Gross CA, Vollmer W. 2015. Coordination of peptidoglycan synthesis and outer membrane constriction during Escherichia coli cell division. Elife 4.

30. Morgan JK, Ortiz JA, Riordan JT. 2014. The role for TolA in enterohemorrhagic Escherichia coli pathogenesis and virulence gene transcription. Microb Pathog 77:42–52.

31. Maltby R, Leatham-Jensen MP, Gibson T, Cohen PS, Conway T. 2013. Nutritional basis for colonization resistance by human commensal Escherichia coli strains HS and Nissle 1917 against E. coli O157:H7 in the mouse intestine. PLoS One 8:e53957.

32. Que JU, Hentges DJ. 1985. Effect of streptomycin administration on colonization resistance to Salmonella typhimurium in mice. Infect Immun 48:169–74.

33. Cordonnier C, Le Bihan G, Emond-Rheault JG, Garrivier A, Harel J, Jubelin G. 2016. Vitamin B12 Uptake by the Gut Commensal Bacteria Bacteroides thetaiotaomicron Limits the Production of Shiga Toxin by Enterohemorrhagic Escherichia coli. Toxins (Basel) 8.

34. Goswami K, Chen C, Xiaoli L, Eaton KA, Dudley EG. 2015. Coculture of Escherichia coli O157:H7 with a Nonpathogenic E. coli Strain Increases Toxin Production and Virulence in a Germfree Mouse Model. Infect Immun 83:4185–93.

35. Curtis MM, Hu Z, Klimko C, Narayanan S, Deberardinis R, Sperandio V. 2014. The gut commensal Bacteroides thetaiotaomicron exacerbates enteric infection through modification of the metabolic landscape. Cell Host Microbe 16:759–69.

36. Zac Stephens W, Burns AR, Stagaman K, Wong S, Rawls JF, Guillemin K, Bohannan BJ. 2016. The 639 composition of the zebrafish intestinal microbial community varies across development. ISME J 10:644–54.

37. Roeselers G, Mittge EK, Stephens WZ, Parichy DM, Cavanaugh CM, Guillemin K, Rawls JF. 2011. Evidence for a core gut microbiota in the zebrafish. ISME J 5:1595–608.

38. Schmidt CE, Shringi S, Besser TE. 2016. Protozoan Predation of Escherichia coli O157:H7 Is Unaffected by the Carriage of Shiga Toxin-Encoding Bacteriophages. PLoS One 11:e0147270.

39. Mast SO. 1947. The food-vacuole in Paramecium. Biol Bull 92:31–72.

40. Kang G, Pulimood AB, Koshi R, Hull A, Acheson D, Rajan P, Keusch GT, Mathan VI, Mathan MM. 2001. A monkey model for enterohemorrhagic Escherichia coli infection. J Infect Dis 184:206–10.

41. Tzipori S, Gibson R, Montanaro J. 1989. Nature and distribution of mucosal lesions associated with enteropathogenic and enterohemorrhagic Escherichia coli in piglets and the role of plasmid-mediated factors. Infect Immun 57:1142–50.

42. Miyamoto Y, Iimura M, Kaper JB, Torres AG, Kagnoff MF. 2006. Role of Shiga toxin versus H7 flagellin in enterohaemorrhagic Escherichia coli signalling of human colon epithelium in vivo. Cell Microbiol 8:869–79.

43. Berin MC, Darfeuille-Michaud A, Egan LJ, Miyamoto Y, Kagnoff MF. 2002. Role of EHEC O157:H7 655 virulence factors in the activation of intestinal epithelial cell NF-kappaB and MAP kinase pathways and the upregulated expression of interleukin 8. Cell Microbiol 4:635–48.

44. Lebeis SL, Bommarius B, Parkos CA, Sherman MA, Kalman D. 2007. TLR signaling mediated by MyD88 is required for a protective innate immune response by neutrophils to Citrobacter rodentium. J Immunol 179:566–77.

45. Meeker ND, Trede NS. 2008. Immunology and zebrafish: spawning new models of human disease. Dev Comp Immunol 32:745–57.

46. Meijer AH, Spaink HP. 2011. Host-pathogen interactions made transparent with the zebrafish model. Curr Drug Targets 12:1000–17.

47. Alsharif G, Ahmad S, Islam MS, Shah R, Busby SJ, Krachler AM. 2015. Host attachment and fluid shear are integrated into a mechanical signal regulating virulence in Escherichia coli O157:H7. Proc Natl Acad Sci U S A 112:5503–8.

48. Leverton LQ, Kaper JB. 2005. Temporal expression of enteropathogenic Escherichia coli virulence genes in an in vitro model of infection. Infect Immun 73:1034–43.

49. Lee DJ, Bingle LE, Heurlier K, Pallen MJ, Penn CW, Busby SJ, Hobman JL. 2009. Gene doctoring: amethod for recombineering in laboratory and pathogenic Escherichia coli strains. BMC Microbiol 9:252.

50. Campellone KG, Robbins D, Leong JM. 2004. EspFU is a translocated EHEC effector that interacts with Tir and N-WASP and promotes Nck-independent actin assembly. Dev Cell 7:217–28.

51. Campellone KG, Giese A, Tipper DJ, Leong JM. 2002. A tyrosine-phosphorylated 12-amino-acid sequence of enteropathogenic Escherichia coli Tir binds the host adaptor protein Nck and is required for Nck localization to actin pedestals. Mol Microbiol 43:1227–41.

52. (ed). 2000. The Zebrafish Book. University Press Oregon, Eugene, OR. Accessed

